# Centrosome identity drives asymmetric microtubule-dependent pushing forces, spindle positioning and daughter cell size

**DOI:** 10.64898/2026.11.12.766129

**Authors:** Imane Naouadir, Gregory Eot-Houllier, Jacques Pécréaux, Hélène Bouvrais, Aude Pascal, Régis Giet

**Affiliations:** Univ Rennes, CNRS, IGDR (Institut de Génétique et Développement de Rennes) UMR 6290; Inserm U1305 Génétique du Cancer

**Keywords:** Astral microtubules, mitotic spindle positioning, centrosome identity, asymmetric pushing force, neural stem cells, asymmetric cell division

## Abstract

Mitotic spindle positioning is essential for tissue architecture and organogenesis. In *Drosophila* neural stem cells, spindle positioning is attributed to apically enriched Dynein generating pulling forces on astral microtubules (MTs). Surprisingly, however, we find that the mitotic spindle consistently adopts a basal position along the apico-basal polarity axis, challenging the canonical model. We show that centrosomes at opposite spindle poles retain an intrinsic asymmetry in MT nucleation capacity inherited from their mother–daughter identity, with the apical daughter centrosome nucleating more MTs than the basal mother centrosome. Equalizing or reversing this asymmetry alters spindle positioning, demonstrating that basal spindle positioning critically depends on asymmetric centrosomal MT nucleation. Centrosome photoablation reveals that astral MTs generate pushing forces against the cell cortex that drive basal spindle positioning, whereas reducing apical Dynein-dependent pulling exacerbates basal spindle displacement, revealing an unexpected role for cortical pulling in limiting spindle displacement. Consistently, 3D Cytosim simulations incorporating cortical pulling forces together with MT nucleation-driven pushing forces, either asymmetric (wild-type or inverted) or symmetric, reproduce the observed spindle positioning. Together, our findings reveal that centrosome identity establishes an intrinsic asymmetry in MT nucleation that converts astral MT pushing into a dominant directional force for spindle positioning. This mechanism counteracts cortical pulling and contributes to daughter cell size asymmetry, uncovering an unexpected role for centrosome identity in determining where cells divide.

## Introduction

Cell division orientation stands as a fundamental pillar in vertebrate tissue growth and homeostasis (di Pietro et al., 2016; Lechler and Mapelli, 2021; McDougall et al., 2019; Sallé and Minc, 2022; Williams and Fuchs, 2013). Seminal studies utilizing *C. elegans* embryos and *Drosophila* Neural Stem Cells (NSCs) have proven instrumental in unraveling the mechanisms governing spindle positioning and cell division orientation (di Pietro et al., 2016; Lechler and Mapelli, 2021; McNally, 2013). In *Drosophila* mitotic NSCs, the spindle orientation machinery and cell fate determinants are anchored at the apical and basal cortex, respectively. Proper alignment of the spindle along the axis of cell polarity regulates the asymmetric distribution of these molecules, influencing the daughter cell’s binary decision to become a NSC or a Ganglion Mother Cell (GMC), destined for differentiation into neurons or glial cells. Failure to orient the spindle along the apico-basal axis results in NSCs expansion at the expense of differentiated cells (Gallaud et al., 2017; Homem and Knoblich, 2012).

Mitotic entry in NSCs is distinguished by a cell-autonomous polarization, marked by the recruitment of the PAR complex (PAR3/Bazooka, PAR6, and aPKC), defining the apical cortex. The Inscuteable protein links the PAR complex to a canonical spindle orientation machinery. This machinery comprises Gαi, Mud and Pins (the homologues of human NuMA and LGN, respectively), together with Dynein, a minus-end-directed MT motor that captures the plus ends of astral MTs and pulls on them (Gallaud et al., 2017). The spindle orientation machinery, asymmetrically localized at the apical cortex, is thought to promote MT catastrophe, associated with Dynein-mediated movement towards centrosome, could generate pulling forces (Laan et al., 2012).

Centrosomes and their astral MTs play a pivotal role in spindle orientation in *Drosophila* (Basto et al., 2006; Megraw et al., 2001). During interphase in NSCs, centrosomes also exhibit functional asymmetry, with the daughter centrosome nucleating more MTs than the mother centrosome (Conduit and Raff, 2010; Rebollo et al., 2007; Rusan and Peifer, 2007). This asymmetry is established through the differential recruitment of Centrobin, that controls the levels of pericentriolar material, and daughter centrosome’s MT nucleation potential (Januschke et al., 2013; Lerit and Rusan, 2013; Zou et al., 2005). Whether that asymmetric MT nucleation is retained during mitosis and contributes to spindle positioning has not yet been investigated.

The pulling canonical spindle orientation machinery, is broadly present from yeast to human cells, including drosophila and nematode (di Pietro et al., 2016; Lechler and Mapelli, 2021; McNally, 2013). MTs can also generate pushing forces by growing against the cell periphery or organelles, which may contribute to spindle positioning or organelle displacements.

Indeed, few examples of MT-driven pushing forces have been described *in vitro* or *in vivo* (Daga et al., 2006; Dogterom and Yurke, 1997; Faivre-Moskalenko and Dogterom, 2002; Garzon-Coral et al., 2016; Holy et al., 1997; Hotani and Miyamoto, 1990; Meaders et al., 2020; Pécréaux et al., 2016; Tissot et al., 2017; Tolić-Nørrelykke et al., 2004; Tran et al., 2001; Zhao et al., 2012; Zhu et al., 2010).

More recently, experimental studies and computational modeling have suggested that both pulling and pushing forces contribute to mitotic spindle positioning and orientation in the one-cell *C. elegans* embryo and other cell types, including *Drosophila* neuroblasts (Bouvrais et al., 2021; Pécréaux et al., 2016; Yeltokov et al., 2026). However, the contribution of astral MT pushing against the cortex to spindle positioning remains poorly understood and has not been directly demonstrated in asymmetrically dividing cells of higher eukaryotes. Here, using asymmetrically dividing *Drosophila* NSCs, we show that basal spindle positioning is primarily driven by the higher MT nucleation capacity of the daughter centrosome, highlighting the critical contribution of astral MT-dependent pushing forces to spindle positioning and the control of daughter cell size.

## Results

### The NSC mitotic spindle is asymmetrically positioned toward the basal cortex

The canonical spindle positioning model posits that the localization of the spindle orientation complex mediates astral MT pulling and orients the spindle within mitotic cells. We took advantage of *Drosophila* NSCs in which the spindle orientation complex is asymmetrically localized at the apical cortex to ensure the correct orientation of the spindle along the apico-basal axis (Figure 1A, left). Apical enrichment of the MT pulling machinery would predict a spindle positioned closer to the apical cortex (Figure 1A, right).

**Figure 1.**
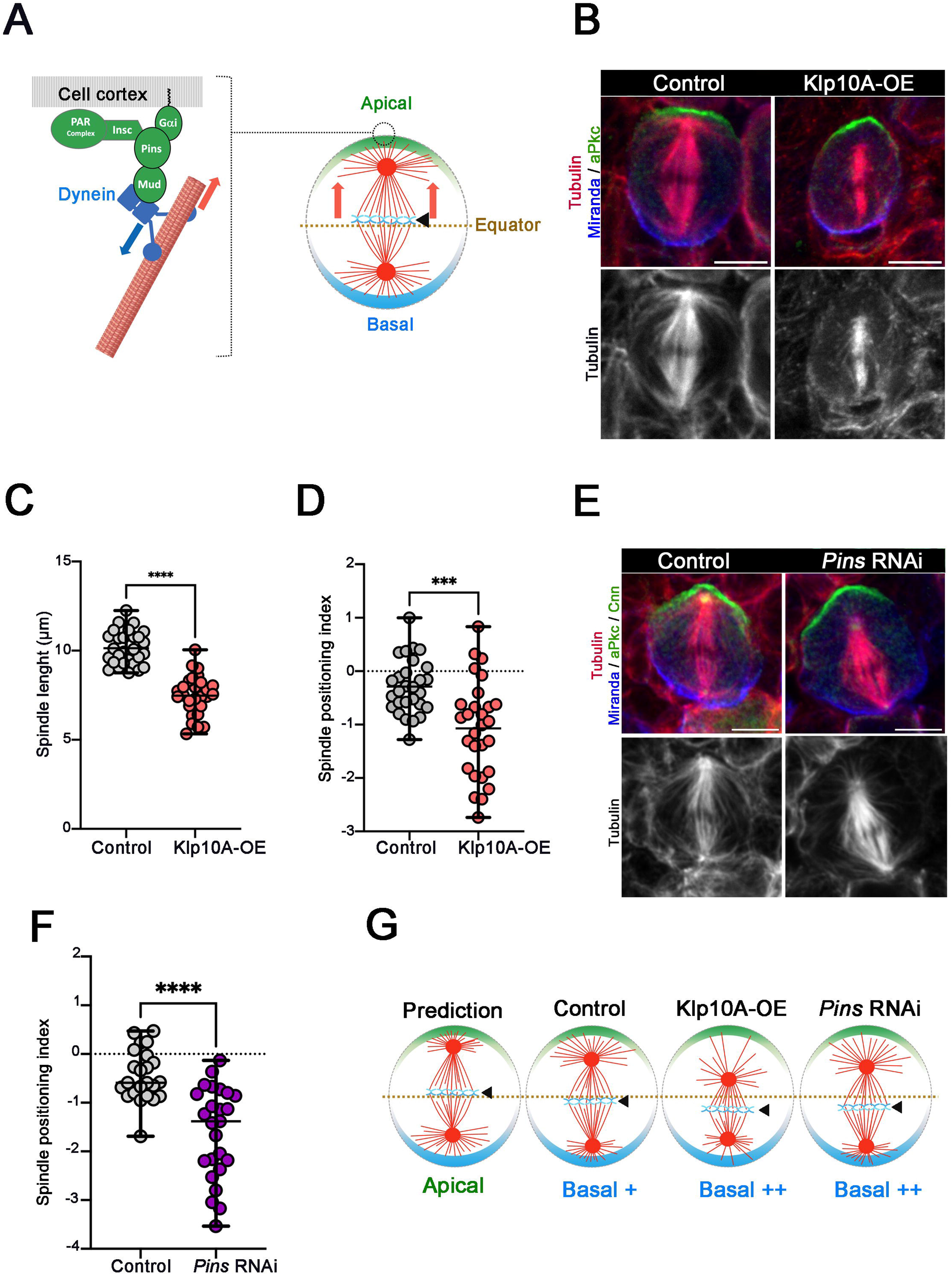
Analysis of spindle positioning in control, Klp10A-OE and *Pins* RNAi NSCs. **A)** Left: Schematic representation of the mitotic spindle orientation machinery in NSCs. The PAR complex (green) is connected via Inscuteable (Insc) to the spindle pulling force generator complex (Gαi, Pins, Mud, and the molecular motor Dynein) at the apical cortex. Cortical dynein (blue) “walks” toward MT minus ends, to generate MT pulling and whole spindle displacement toward the cortex (red arrow). Right: Schematic of a *Drosophila* NSC cell during metaphase. The spindle orientation machinery is located at the apical cortex (green crescent) and defines a cortical region where astral MT pulling events occur to orient the spindle along the apico-basal axis. These pulling forces are expected to position the spindle toward the apical cortex. The NSC equator is displayed as a brown dotted line. The basal cortex, where cell fate determinants reside, is displayed as a blue crescent. **B)** Localization of MTs (red, and lower panels in monochrome), aPKc (green, apical cortex) and Miranda (blue, basal cortex) in control (Left) and Klp10A overexpressing cells (KLP10A-OE, right). Scale bar is 5 µm. **C)** Dot plot showing the mean (± sd) mitotic spindle length in control (10.14 µm ± 0.95, *n =* 28) and Klp10A-OE metaphase NSCs (7.48 µm ± 1.11, *n =* 28). ****, *P* ≤ 0.001 (Mann-Whitney test). **D)** Spindle positioning index analysis in control or Klp10-OE mitotic cells. (See also Figure S1 and Methods). The mitotic spindles are positioned basally in control (−0.30 ± 0.51, *n =* 28) and Klp10A-OE (−1.07 ± 0.89, *n =* 28) NSCs. ***, *P* ≤ 0.001(Mann-Whitney test). **E)** Control, and *Pins* RNAi NSCs during metaphase. aPKC (apical cortex) and Cnn are displayed in green. MTs are shown in red (and lower panels in monochrome) and Miranda (basal cortex) in blue. Scale bar is 5 µm. **F)** Spindle positioning index analysis (± sd) in control or *Pins* RNAi mitotic cells. The mitotic spindles are positioned basally in control (−0.51 ± 0.61, *n =* 24) and *Pins* RNAi (−1.61 ± 0.94, *n =* 25) NSCs. ****, *P* ≤ 0.0001. (Mann-Whitney test). **G)** Scheme representing spindle position when driven by an apical only Dynein pulling mechanism (prediction). versus measured positioning in control, Klp10A-OE cells and *Pins* RNAi mitotic NSCs.

To assess the position of the mitotic spindle along the apico-basal axis during metaphase in Drosophila larval NSCs, we measured the distances between each centrosome and the cell cortex and used these values to calculate a spindle positioning index indicating whether the spindle was positioned toward the apical or basal cortex (Figure 1B, S1). In parallel to controls, we also elevated the levels of the *Drosophila* Kinesin 13 (Klp10A), a MT-depolymerizing kinesin to decrease the number of polymerizing astral MTs and shorten the spindles as previously described (Figure 1B and 1C) (Goshima and Vale, 2005; Mennella et al., 2005; Morales-Mulia and Scholey, 2005; Thomas et al., 2021). We found that the majority of control NSCs displayed basally-positioned mitotic spindle (Figure 1B and 1D). Moreover, basal spindle positioning was also present in Klp10A-OE NSCs (Figure 1D). Altogether, these data show that, by contrast to the prediction (Figure 1G, left panel), NSCs mitotic spindles are positioned toward the basal cortex, opposite to the pulling machinery (Figure 1G, middle panels). This suggests that additional mechanisms, other than MT pulling, contribute to adjust spindle position toward the basal cortex along the polarity axis, independent of spindle size (Figure 1G, right panel).

### NSC asymmetric basal positioning is counterbalanced by cortical pulling forces mediated by the Dynein adaptor Pins

The abundant literature has shown that Dynein motors, targeted at the apical cortex via adaptor proteins, are instrumental for orienting the spindle by pulling on astral MTs (Figure 1A) (di Pietro et al., 2016; Kiyomitsu and Boerner, 2021; Lechler and Mapelli, 2021; McNally, 2013; Nakajima, 2018; Tarannum et al., 2022; Zellag et al., 2025). However, the basal spindle axial positioning observed in figure 1D was not in agreement with Dynein-mediated MT pulling playing a major role in that process. We therefore examined in detail the position of the mitotic spindle with diminished Dynein levels at the cell cortex upon knock down of its cortical adaptor Pins. Earlier studies showed that *pins* null mutants displayed severe spindle orientation defects precluding positioning analyses along the apico-basal axis (Izumi et al., 2006; Siegrist and Doe, 2005; Siller et al., 2006). We therefore measured spindle positioning under conditions where spindle orientation was maintained, by partially depleting Pins by RNAi (Figure 1E). *Pins* RNAi cells exhibited enhanced spindle basal positioning compared to control NSCs (Figure 1E and 1F). This result suggests that the spindle orientation machinery equilibrates a basal spindle positioning mechanism (Figure 1G, right panel).

### NSC centrosomes retain an asymmetric MT nucleation associated with their identity

We then wondered if astral MTs, nucleated by the centrosomes at spindle poles could play a role in basal spindle positioning. During interphase, *Drosophila* NSCs mother/basal and daughter/apical centrosomes exhibit strong differences in their composition and nucleation potential. The daughter centrosome harbors pericentriolar material (PCM), nucleates MTs and remains associated with the apical cortex. By contrast, the mother centrosome loses its PCM, MTs and migrates toward the basal cortex (Conduit et al., 2010; Izumi et al., 2006; Januschke and Gonzalez, 2010; Januschke et al., 2011; Rebollo et al., 2007; Rusan and Peifer, 2007). We therefore examined if centrosome asymmetry was retained during cell division in NSCs (Figure S2). When monitoring the PCM component Centrosomin (Cnn), by quantifying the fluorescence intensity of endogenously sfGFP-tagged Cnn (Mukherjee et al., 2020), we confirmed previous results: at nuclear envelope breakdown (NEBD), the daughter centrosome exhibited high levels of Cnn-sfGFP and a large MT aster (Figure S2A, green triangle and Video S1). The mother centrosome started to be detected when it incorporated Cnn-sfGFP few minutes before mitosis (Figure S2A, blue triangle and Video S1). Strikingly, during cell division, from NEBD to telophase, this asymmetry in Cnn-sfGFP levels between the two centrosomes was conserved (Figure S2B and Video S1). Analyses of Cnn-sfGFP fluorescence ratio between both centrosomes revealed that the apical/daughter centrosome contains in average 2 to 3 times more Cnn-sfGFP than the basal/mother centrosome during mitosis (Figure S2C). Finally, we quantified if this asymmetric PCM composition was correlated with asymmetric MT nucleation. Analysis of the number of *de novo* nucleated MTs, measured during prometaphase and metaphase using an endogenously tagged Eb1-tdGFP protein (Sun et al., 2021) showed that the apical/daughter centrosome spindle pole nucleated in average 1.5 time more MTs toward the cell cortex than the daughter/basal centrosome (Figure S3 A and B and Video S2). Altogether, this indicates that both apical and basal centrosomes retain asymmetric levels of Cnn-sfGFP, and asymmetric MT nucleation potential during mitosis, with this asymmetry correlating with their mother/daughter interphase origin.

### Loss of asymmetric MT nucleation prevents basal spindle positioning

We thus hypothesized that the centrosome identities, associated with their asymmetric MT nucleation could be responsible for the observed biased positioning of NSC spindles toward the basal cortex. We therefore aimed to generate spindle poles harboring centrosomes, with similarly high or low MT nucleation capacity to assess their impact on spindle positioning (Figure 2A). In NSCs, pioneering work has shown that centrosome identity and their asymmetric MT nucleation are controlled by Centrobin (Cnb) levels (Figure 2A). To convert both centrosomes into “daughter-like” centrosomes with high MT nucleation capacity, we targeted Cnb to both centrosomes using a version of the protein tagged with a PACT/centrosome targeting sequence (Januschke et al., 2013; Lerit and Rusan, 2013) (Figure 2A top). In parallel, we converted both centrosomes into mother-like centrosomes with low MT nucleation capacity by RNAi-mediated depletion of Cnb (Januschke et al., 2013) (Figure 2A bottom). Using immunostaining analyses of endogenous Cnn, we assessed its level at the two centrosomes in the different conditions. In the control, the daughter centrosome exhibited higher Cnn levels than the mother one during metaphase (Figure 2B left, Figure 2C and Figure S2D). The centrosomes of mitotic Cnb-PACT mitotic NSCs exhibited similar “high” amount of Cnn proteins indicating that both centrosomes harbored a daughter-like behavior (Figure 2B middle, Figure 2C and Figure S2D). *Cnb* RNAi resulted in mitotic spindles in which both centrosomes carried low, equivalent levels of Cnn, a “mother-like” behavior although Cnn levels were lower than those of mother centrosomes in control NSCs. (Figure 2B right, Figure 2C and Figure S2D). Under these symmetric centrosome MT nucleation conditions, *Cnb* RNAi or Cnb-PACT triggered the loss the basal spindle positioning observed in control metaphase NSCs (Figure 2D). Cnb-PACT spindles with symmetric high-nucleating centrosomes became equatorial or slightly apical, while the majority of *Cnb* RNAi low-nucleating centrosomes were preferentially positioned toward the apical cortex. Thus, the conversion of asymmetric centrosomes into symmetric centrosomes harboring similar MT nucleation potential is sufficient to change the basal position of the mitotic spindle toward an apical position (Figure 2E).

**Figure 2.**
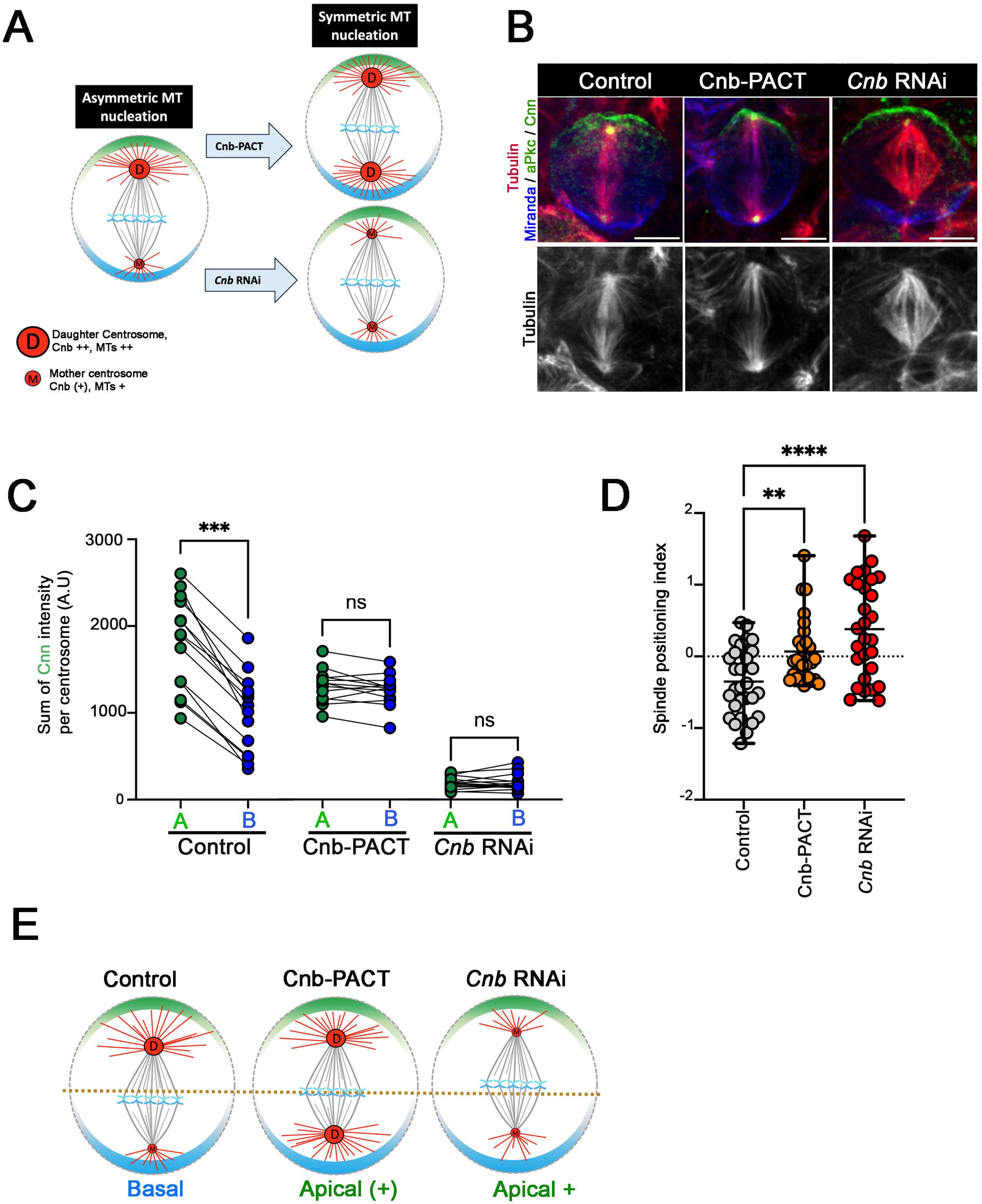
Analysis of mitotic spindle axial positioning in NSC with symmetric centrosomes. **A)** Strategy to generate centrosomes with symmetric low or high MT-nucleating activity. Centrobin (Cnb) levels regulate the recruitment of peri-centriolar material, and asymmetric MT nucleation. Targeting Cnb to both centrosomes using a PACT-domain (Cnb-PACT) or lowering Cnb levels (*Cnb* RNAi) produces symmetric daughter-like/high nucleating or mother-like/low MT nucleating centrosomes, respectively. **B)** Representative images of control, Cnb-PACT, and *Cnb* RNAi NSCs during metaphase. aPKC marks the apical cortex and Cnn labels centrosomes (green), MTs are shown in red (and lower panels in monochrome) and Miranda labels the basal cortex (blue). Scale bar: 5 µm. **C)** Quantification of the average intensity (± sd) of endogenous Cnn at the apical (green) and basal (blue) poles of the spindle during metaphase in control apical: 1806.4 ± 539.4, *n =* 14; basal: 986.0 ± 452.9, *n =* 14), Cnb-PACT (apical: 1302.0 ± 193.2, *n =* 13; basal: 1253.0 ± 183.6, *n =* 13), and *Cnb* RNAi (apical: 182.5 ± 68.64, *n =* 16; basal: 193.3 ± 92.75, *n =* 16) NSCs. ns, not significant; ***, *P* ≤ 0.001 (Mann–Whitney test). The daughter and the corresponding mother centrosome for each cell are connected by straight lines. **D)** Spindle positioning index analysis (± sd) of control, Cnb-PACT and *Cnb* RNAi NSCs during metaphase. The mitotic spindles are positioned basally in control cells (−0.35 ± 0.47, *n =* 29), Cnb-PACT (0.07 ± 0.44, *n =* 28) and *Cnb* RNAi NSCs (0.38 ± 0.68, *n* = 28). **, *P* ≤ 0.01; ****, *P* ≤ 0.0001. (Mann-Whitney test). **E)** Scheme recapitulating the axial spindle position in control and in mitotic NSCs harboring symmetric high-nucleating (Cnb-PACT) or symmetric low-nucleating (*Cnb* RNAi) centrosomes.

### Inverting mother/daughter centrosome configuration positions the spindle toward the apical cortex

The centrosome conversion experiments into symmetric centrosomes suggested that centrosome identity and associated asymmetric MT nucleation were crucial for spindle positioning in NSCs. We hypothesized that the basal spindle positioning observed in control NSCs was dependent upon the larger number of astral MTs nucleated by the daughter centrosome located at the apical cortex vicinity. To challenge this hypothesis, we decided to invert the position of the daughter/high nucleating and mother/low nucleating centrosomes along the apico-basal axis following interference with the centrosome separation. Centrosome separation occurs during interphase in NSCs. The mother centrosome, deprived of MTs is transported along astral MTs nucleated by the daughter centrosome. This transport is performed by Kinesin-1 toward the basal cortex, resulting in the mother centrosome being basally positioned when mitosis starts (Figure 3A, top). Mutations in the Kinesin-1 heavy (Khc) chain prevent the mother centrosome migration and frequently leads to an inverted centrosome configuration relative to the polarity axis in metaphase cells (Gallaud et al., 2014; Hannaford et al., 2022) (Figure 3A, bottom). Analysis of centrosomal Cnn levels, as a readout of centrosome identity and MT nucleation capacity, showed that in 43% of the *Khc^27^/Khc^63^* metaphase NSCs, the daughter/high-nucleating and the mother/low-nucleating centrosomes were inverted relative to the apico-basal axis (Figure 3A and B). *Khc^27^/Khc^63^* NSCs that harbored a wild-type centrosomes configuration retained a basal spindle position similar to control cells (Figure 3B, left and middle panel, and 3C, left and middle). In contrast, *Khc^27^/Khc^63^* metaphase cells with “inverted” centrosomes harbored spindles that were mostly positioned in an apical position (Figure 3B and 3C, right). Altogether, these results support a model in which the daughter centrosome dictates the axial mitotic spindle position within NSCs, independently of the apical polarity complex and spindle orientation machinery (Figure 3D).

**Figure 3.**
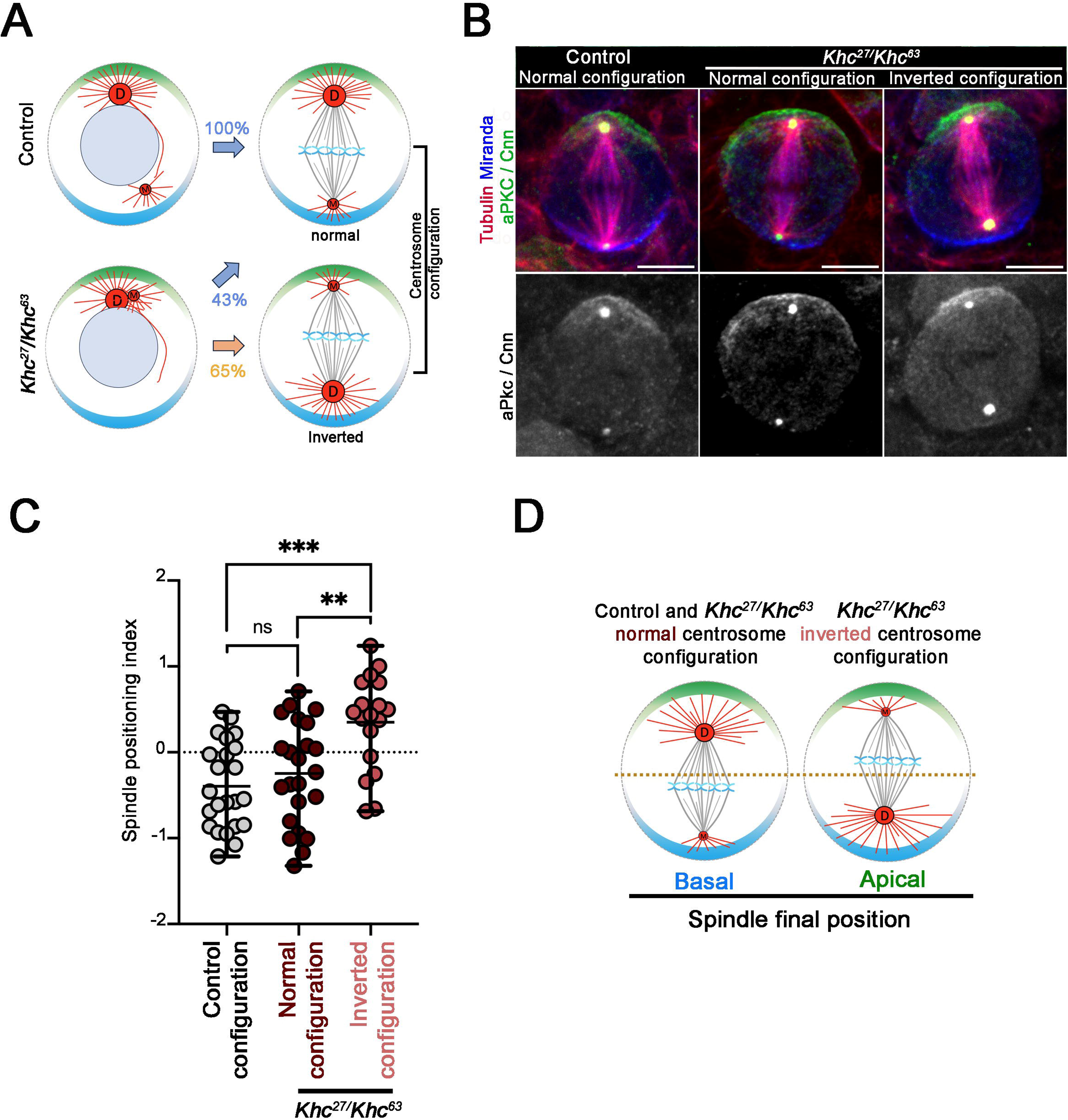
Analysis of mitotic spindle axial positioning in NSC with an inverted centrosome configuration. **A)** Strategy to generate mitotic NSCs with an inverted centrosome configuration. During interphase Kinesin-1 promotes the transport of the MT-deprived mother centriole toward the basal cortex. Before NEBD, this mother centriole recruits PCM to nucleate and organize MTs at the future basal spindle pole. In *Khc^27^Khc^63^* NSCs, centriole separation is impaired and when the mother centrosome nucleates astral MTs prior NEBD, mother centrosome/apical cortex interactions can be produced leading to inverted centrosome configurations in which the mother centrosomes/spindle pole becomes connected to the apical cortex while the daughter centrosome is positioned close to the basal cortex (43%, *n* =18/41). **B)** Representative metaphase in control and *Khc^27^/Khc^63^* NSCs during metaphase. Daughter and mother centrosomes are distinguished by their relative size and Cnn fluorescence intensity, with the daughter centrosome displaying a larger size and stronger Cnn signal than the mother centrosome. The left panel shows a control NSC with normal daughter centrosome positioning at the apical cortex. The middle panel shows a *Khc^27^/Khc^63^* NSC with daughter centrosome positioning similar to the control (wild-type centrosome configuration). The right panel shows a *Khc^27^/Khc^63^* with an inverted spindle and the daughter centrosome positioned at the basal cortex (inverted centrosome configuration). aPKC (apical cortex) and Cnn are displayed in green. MTs are shown in red and Miranda (basal cortex) in blue. Scale bar is 5 μm. **C)** Quantification of the spindle positioning index (± sd) in control and *khc^27^/khc^63^* NSCs during metaphase. In control and *khc^27^/khc^63^* NSCs with wild-type centrosome configuration, spindles are positioned basally (WT: −0.39 ± 0.51, *n* = 23; *khc^27^/khc^63^* normal: −0.25 ± 0.59, *n* = 23). In contrast, *khc^27^/khc^63^*NSCs with inverted centrosome configuration display an axial spindle position toward the apical cortex (0.35 ± 0.55, *n* = 18). ns, not significant, *P* > 0.05; *, *P* ≤ 0.05; **, *P* ≤ 0.01; ***, *P* ≤ 0.001 (One-way ANOVA followed by Tukey’s post hoc multiple comparisons test). **D)** Scheme representing axial spindle position in mitotic NSCs with control or inverted centrosome configurations.

### Centrosome ablation reveals a contribution of astral microtubule pushing forces to spindle positioning

Imaging of astral MT growing plus ends in NSCs revealed many Eb1-tdGFP comets polymerizing against the cell cortex, suggesting that growing MTs were able to exert pushing forces in this context (Video S2). We therefore used laser ablation in live NSCs, for probing microtubule force generation in spindle positioning, and analyzed the resulting spindle displacement (Khatri et al., 2022; Odell et al., 2019; Thomas et al., 2021; Thomas and Giet, 2022; Wu et al., 2024). To determine the spatial contribution of astral MTs nucleated by each centrosome to force generation, we selectively ablated either the daughter (apical) or the mother (basal) centrosome in NSCs, thereby disrupting the associated astral MTs. We then tracked the displacement of the opposite spindle pole (Figure 4A and B). We found that ablation of the basal (mother) centrosome induced apical spindle pole displacement toward the basal cortex (Figure 4A and C). Conversely, ablation of the apical (daughter) centrosome triggered displacement of the opposite basal spindle pole toward the apical cortex along the polarity axis (Figure 4B and D). Thus, ablation of either centrosome causes the opposite spindle pole to move away from its adjacent cell cortex. This behavior supports the idea that MTs growing from both the daughter and mother centrosomes generate pushing forces against the cell cortex that primarily contribute to mitotic spindle positioning, over pulling forces at the apical cortex.

**Figure 4.**
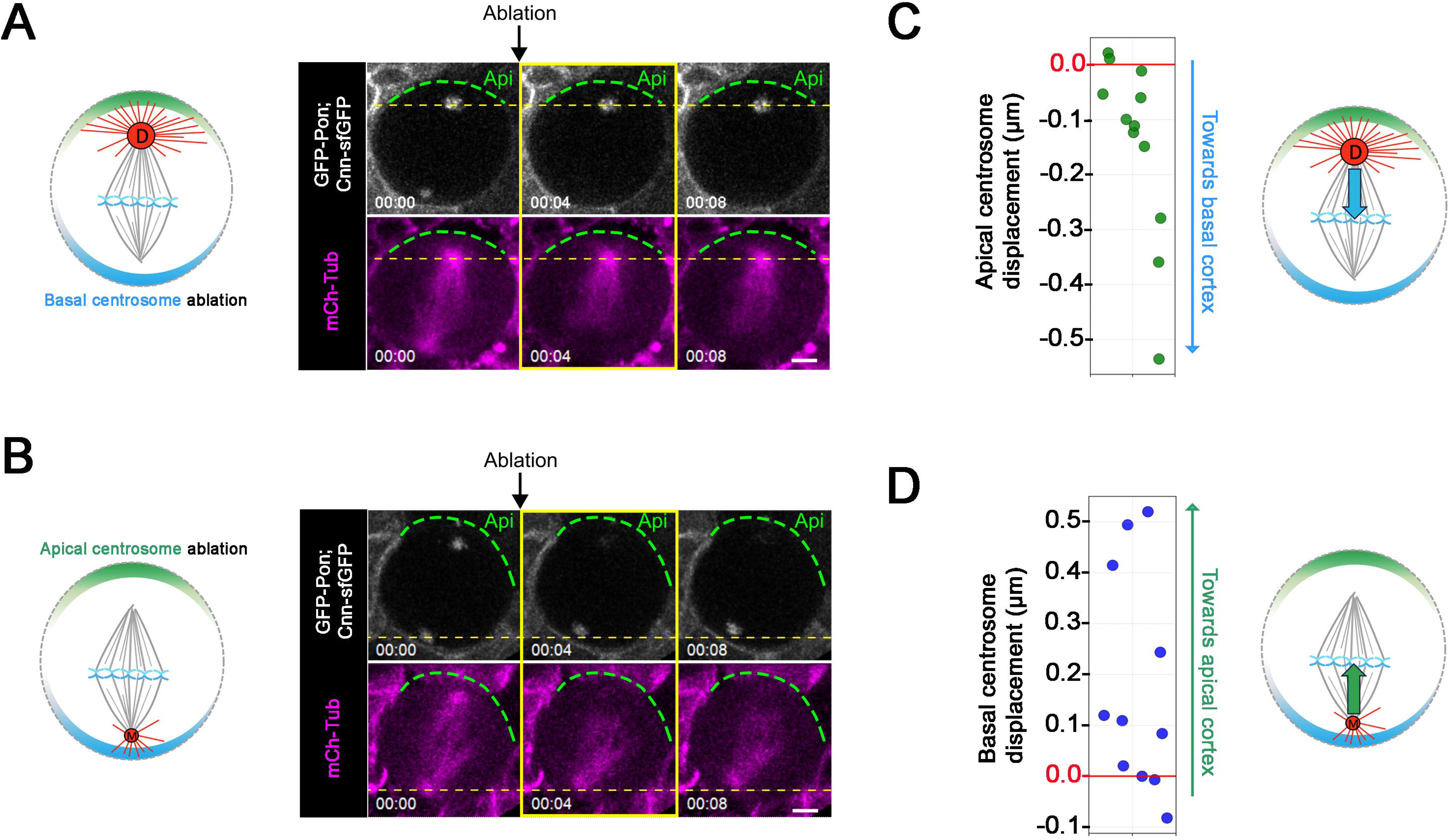
Analysis of centrosome displacement following laser ablation. **A) Left:** Scheme of a NSC with ablated basal/mother centrosome. **Right:** live NSC subjected to mother centrosome ablation. **B) Left:** Scheme of a NSC with ablated apical/daughter centrosome. **Right:** pre-ablation: -4 s and post-ablation: 0 and 4 s. Cnn-GFP and Pon-GFP are shown in green, MTs are shown in purple. The apical cortex is displayed with a dotted green line. In A and B, pre -ablation: 0 s post-ablation: 4 and 8 s and the scale bar is 2.5 μm. **C)** Quantification of the apical daughter centrosome displacement (μm) following ablation of the basal mother centrosome (left). The right scheme summarizes the basal displacement of the daughter following ablation of the mother centrosome. **D)** Quantification of basal/mother centrosome displacement (μm) following ablation of the mother/apical centrosome. The right scheme summarizes the basal displacement of the mother following ablation of the daughter centrosome. The zero is indicated by a red line that sets the initial position of the unablated centrosome before ablation of its opposite counterpart. Negative and positive values indicate displacement toward the basal cortex and apical cortex, respectively.

### Simulation recapitulates spindle positioning dependence on centrosomal nucleation capacities

To further test the importance of asymmetric cortical pushing force in *Drosophila* NSC spindle positioning, we constructed a simplified computational model using Cytosim, a well-established simulation software for cytoskeletal-based cellular processes (Nedelec and Foethke, 2007). Cytosim has previously been used to investigate MT-organizing center positioning (Boudreau et al., 2019; Cueff et al., 2026; Kozlowski et al., 2007). To mimic wild-type spindle positioning, we included asymmetric nucleation of astral MTs at the centrosomes, based on experimental counting from confocal microscopy (Figure S3). We simplified cortical pulling forces to a constant force exerted on the apical pole, benefiting from our additions to the core Cytosim framework (Pecreaux, 2024). Astral MTs nucleated from the centrosomes were simulated as dynamic filaments capable of generating cortical pushing forces by growing against the cell cortex. At the start of the simulation, the mitotic spindle (with a length equal to the average metaphase spindle; Figure 1C, left) was positioned at the cell center. We then ran the simulation for 5 min, including 30 s for initial relaxation. Confirming that the simulation well capture the dynamics, we observed oscillations of the mitotic spindle orientation as previously reported *in vivo*, (Siller and Doe, 2008). For this static analysis, we considered that the final position of the spindle, reached after 3 min, similarly to its *in vivo* counterpart. Furthermore, we averaged the spindle position over the last two minutes to assess displacement along the apico-basal axis at the end of the simulation at a time-scale beyond oscillations (Video S1). Running 100 simulations that mimic wild-type divisions, with the nucleation capacity set in agreement with experimental measurements (Figure S3), we revealed an overall basal spindle final position (Figure 5A and B left, Video S3), consistent with *in vivo* observations (Figure 1B & D, left). This suggests that a pushing force imbalance, resulting from the asymmetric nucleation capacity of the centrosomes, higher on the apical side, is sufficient to recapitulate wild-type spindle positioning in the presence of apical pulling forces. To challenge the key contribution of centrosome nucleation capacity, we simulated three centrosome-nucleation perturbations that were studied experimentally: high and symmetric nucleation (mimicking Cnb-PACT, Figure 2A), low and symmetric nucleation (mimicking *Cnb* RNAi, Figure 2A), and inverted asymmetric nucleation (mimicking *khc^27^/khc^63^*, Figure 3A). The inverted asymmetric nucleation led to apical spindle positioning (Figure 5A’’’and B right, Video S6), phenocopying the *khc^27^/khc^63^* experiment (Figure 3B, C). High and symmetric nucleation caused spindles to be positioned close to the equator (Figure 5A’ and B middle left, Video S4), while low and symmetric nucleation resulted in apical spindle localization (Figure 5A’’ and B middle right, Video S5). These simulations were consistent with *in vivo* observations (Figure 2D). Overall, the simulations confirmed that spindle positioning critically depends on centrosomal nucleation capacity, consistent with cortical pushing forces dominating cortical pulling. As a consistency check, we varied the apical pulling force from 0 to 50 pN while preserving wild-type asymmetry in centrosomal MT nucleation. In these simulations, asymmetric centrosomal MT nucleation alone was sufficient to keep the spindle in a basal position under low apical pulling conditions (Figure S4, Video S7).

**Figure 5.**
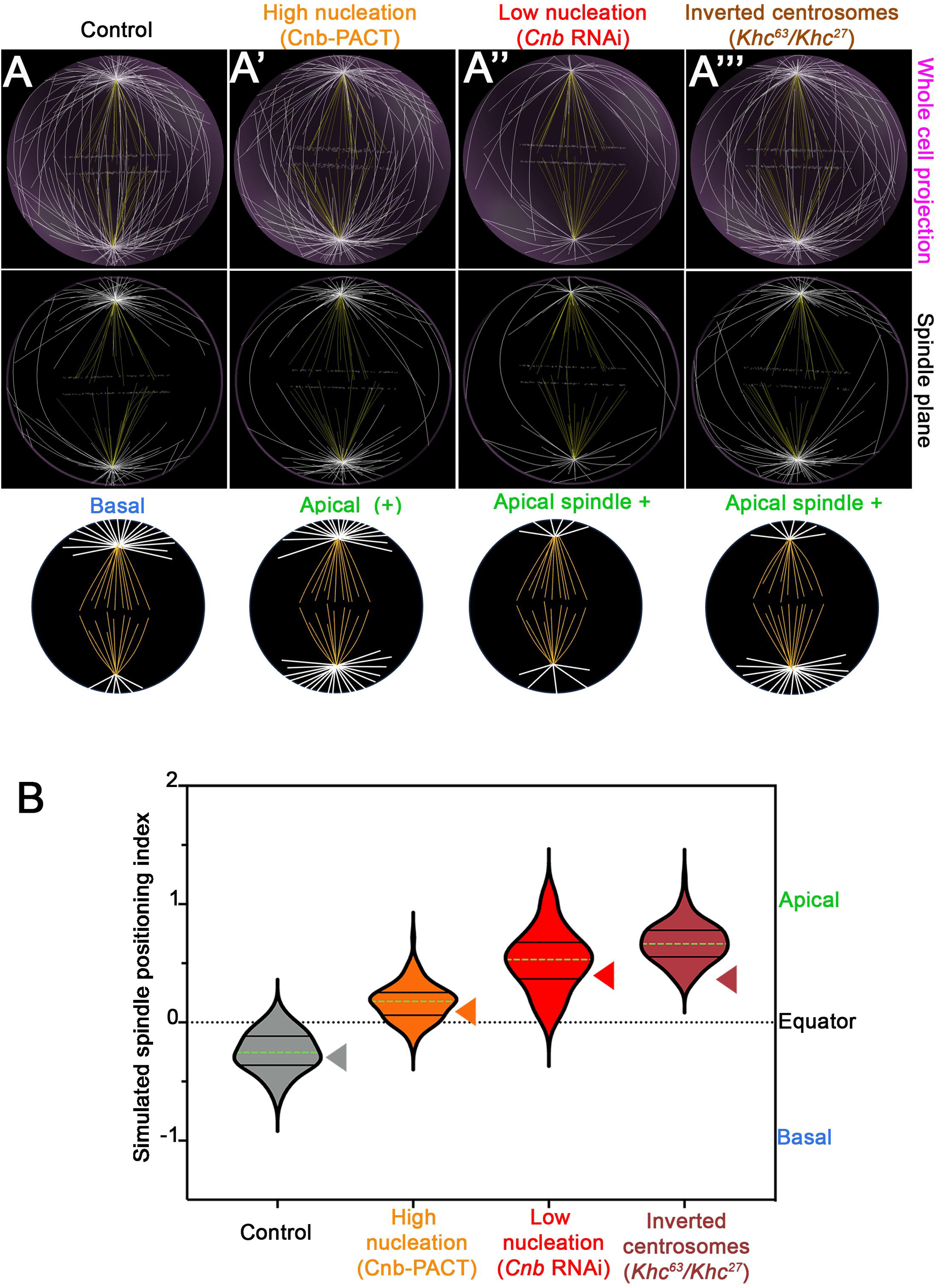
Cytosim simulation of spindle positioning in NSCs under varying centrosomal microtubule nucleation activities A-A’’’) Representative images of the final simulated spindle positions in mitotic NSCs under four conditions: (A) control NSCs with apical MT pulling forces and asymmetric MT nucleation, in which the apical (daughter) centrosome nucleates 1.5-fold more MTs than the basal (mother) centrosome; (A′) NSCs with apical MT pulling forces and symmetric high MT nucleation (mimicking Cnb-PACT); (A′′) NSCs with apical MT pulling forces and symmetric low MT nucleation (mimicking *Cnb* RNAi); and (A′′′) NSCs with apical MT pulling forces and inverted centrosome configuration (mimicking *Khc^27^/Khc63*). First and second rows show 3D view and single optical sections of the simulations, respectively. In the third raw, schemes represent axial spindle positions for the indicated simulated conditions. **B)** Violin plots showing the distribution of the final spindle positioning index along the NSC polarity axis, expressed as log₂(B/A), under the four conditions described in (A). Mean values ± sd are as follows: control (asymmetric nucleation), −0.25 ± 0.17; Cnb-PACT (symmetric high nucleation), 0.17 ± 0.15; *Cnb* RNAi (symmetric low nucleation), 0.52 ± 0.26; and *Khc27/Khc63* (inverted centrosomes), 0.67 ± 0.17. For each simulated condition, *n* = 100. Colored triangles indicate the corresponding experimental measurements from Figures 1–3.

Altogether, these simulations successfully reproduce *in vivo* observations (Figure 2D), further supporting that spindle positioning in NSCs critically depends on centrosomal nucleation capacity and that cortical pushing forces dominate over cortical pulling.

### Perturbation of basal spindle positioning impairs daughter cells sizes

Asymmetrically dividing cells are characterized by an asymmetric cleavage furrow position, which produces daughter cells of different sizes. In NSCs, the final asymmetric position of the furrow components is determined by cortical polarity, and the spindle midzone (Roubinet et al., 2017; Thomas et al., 2021). To assess whether the final position of the cleavage furrow is affected when spindle displacement is impaired, we used live imaging to measure the daughter cell size asymmetry index in control cells and in cells in which the spindle was apically positioned (Figure 6). For this purpose, we used *Khc RNAi*, which retain cortical polarity and centrosome MT nucleation but give rise to two populations of dividing cells: “wild-type configured” cells, in which a highly nucleating centrosome lies at the apical cortex, resulting in a basal spindle position similar to wild-type cells, and “inverted configured” cells, in which a highly nucleating centrosome lies at the basal cortex, resulting in an apical spindle position (Figure 3). We found that *Khc* RNAi cells with a wild-type centrosome configuration exhibited daughter NSC and GMC sizes comparable to those of controls, whereas cells with an inverted centrosome configuration produced larger GMCs and smaller NSCs (Figure 6A–C), thereby reducing the NSC-to-GMC diameter ratio and the extent of daughter cell size asymmetry (Figure 6D). Together, these results support a role for basal spindle positioning, driven by asymmetric MT pushing forces, in maintaining daughter cell size asymmetry.

**Figure 6.**
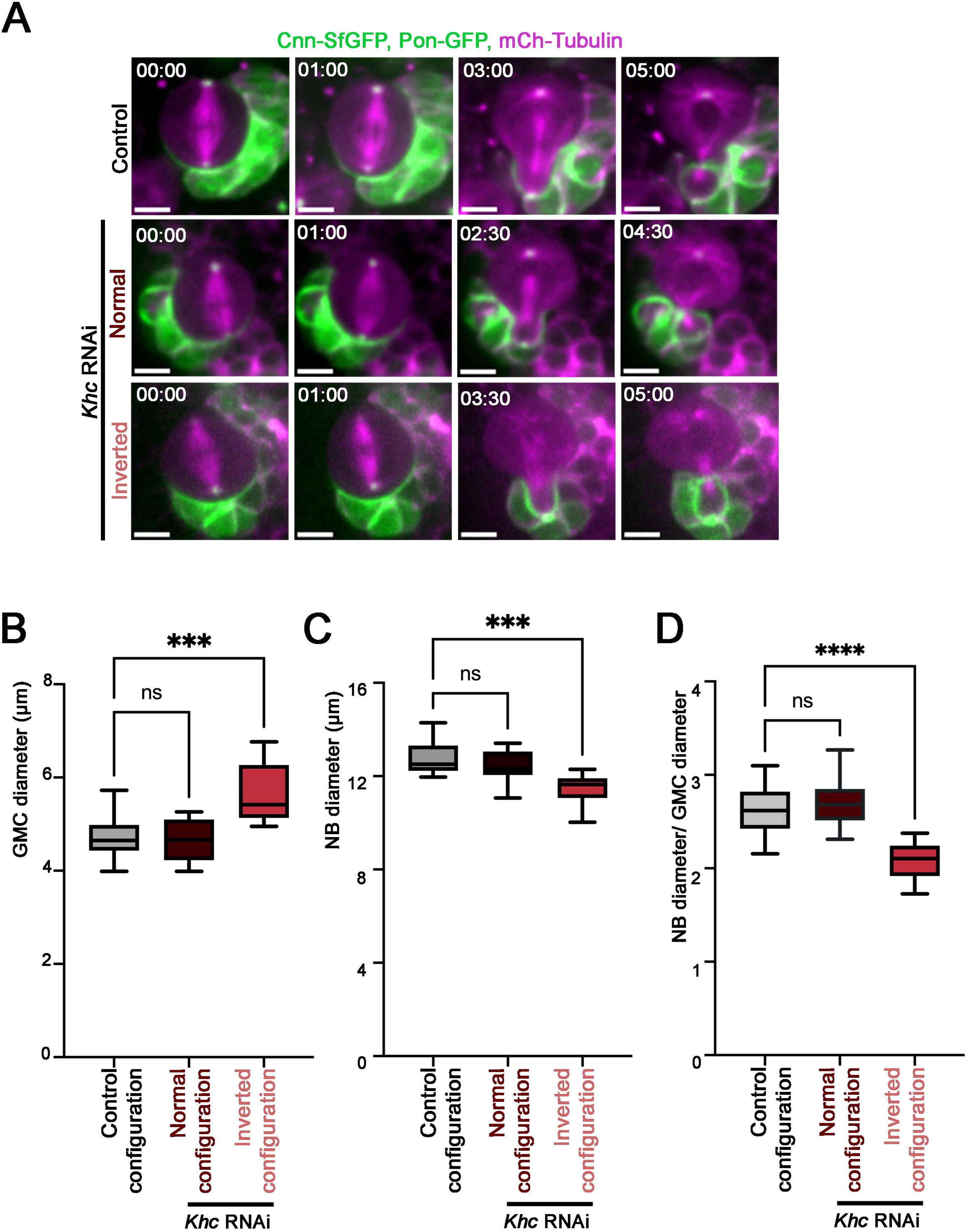
Analysis of cell size asymmetry during cytokinesis in NSCs with inverted centrosome configuration. **A)** Selected frames showing NSCs undergoing cell division in controls (top panels) and *Khc* RNAi cells displaying either normal (middle panels) or inverted centrosome configuration (bottom panels). Cells express mCh-tubulin (purple), Cnn-sfGFP and GFP-Pon-LD (green). Scale bar, 10 µm. Time is indicated in min. B) Box-and-whisker plots showing the mean (± sd) GMC diameter after cell division in control cells (*n* = 12, 4.74 ± 0.50) and *Khc* RNAi cells exhibiting either normal (*n* = 12, 4.64 ± 0.45) or inverted centrosome configuration (*n* = 12, 5.66 ± 0.65). ns, not significant; ***, *P* ≤ 0.001 (Mann-Whitney test). **C)** Box-and-whisker plots showing the mean (± sd) NSC diameter after cell division in control cells (*n* = 12, 12.76 ± 0.78) and *Khc* RNAi cells exhibiting either normal (*n* = 12, 12.45 ± 0.66) or inverted spindle orientation (*n* = 12, 11.48 ± 0.68). ns, not significant; ***, *P* ≤ 0.001 (Mann-Whitney test). **D)** Box-and-whisker plots showing the mean (± sd) NSC-to-GMC diameter ratio after cell division in control cells (*n* = 12, 2.61 ± 0.27) and *Khc* RNAi cells exhibiting either normal (*n* = 12, 2.70 ± 0.26) or inverted spindle orientation (*n* = 12, 2.08 ± 0.20). ns, not significant; ****, *P* ≤ 0.0001 (Mann-Whitney test).

## Discussion

### Spindle positioning in NSC does not follow a dominant Dynein pulling model

NSC mitotic spindle orientation relies on cortically localized Pins that recruits the Dynein adapter Mud at the apical cortex (di Pietro et al., 2016; Kiyomitsu and Boerner, 2021; Lechler and Mapelli, 2021; McNally, 2013; Nakajima, 2018; Tarannum et al., 2022; Zellag et al., 2025). The Dynein motor activity in turns generates the minus-end-directed forces that are instrumental to orient the spindle along the polarity axis. Therefore, when Dynein apical recruitment is abolished, the spindle fails to align along the apico-basal axis, which impairs asymmetric segregation of cell fate determinants into the daughter cells and compromises brain homeostasis (Bowman et al., 2006; Izumi et al., 2006; Siller et al., 2006).

Yet Dynein-mediated MT pulling alone cannot account for the full range of spindle positioning and orientation phenotypes. In several model systems, spindles even align orthogonally to cortical domains enriched in Dynein adaptors (Neville et al., 2023; Saleh et al., 2023).

NSCs are particularly informative in this respect because of their intrinsic asymmetry. First, the cortical MT-pulling machinery is restricted to the apical cortex. Second, the two centrosomes differ in their MT-nucleating capacity, and the more active daughter centrosome is retained close to the apical cortex, where the canonical pulling machinery sits. Both asymmetries predict that the greater number of astral MTs and the enrichment of Dynein at the apical side should draw the spindle toward the apical cortex. Strikingly, we observe the opposite: at metaphase, spindles are consistently positioned basally. Apical pulling therefore sets orientation of the spindle but cannot be the sole determinant of spindle position. Additional regulatory mechanisms must be at work.

### Spindle positioning relies on the asymmetric MT nucleation potential of centrosomes inherited by their mother or daughter origins

Spindle orientation in NSCs is dependent upon the presence of astral MTs nucleated by the centrosomes in NSCs (Basto et al., 2006). Our data now show that centrosomes identity regulates the basal positioning of the spindle along the apico-basal axis. In NSCs, the two centrioles split immediately after cytokinesis, the daughter centriole harbors PCM, nucleates MTs and is retained at the vicinity of the apical cortex. The mother centriole loses its PCM, is deprived of MTs and transported by Khc toward the basal cortex along the astral MTs nucleated by the daughter centriole (Conduit and Raff, 2010; Gallaud et al., 2014; Januschke et al., 2011; Rebollo et al., 2007; Rusan and Peifer, 2007). This migrating mother centriole only starts to recruit PCM before NEBD, to assemble the basal spindle pole. Interestingly, we show that this MT nucleation asymmetry is maintained during mitosis, with the apical daughter centrosome harboring more PCM and nucleating, on average, 50% more MTs than the basal mother centrosome. Abolishing this asymmetry by generating centrosomes with either uniformly low or high MT nucleation capacity resulted in a more apical spindle position. However, this apical displacement was more pronounced in cells with low MT numbers than in those with high MT numbers. We therefore hypothesize that apical pulling relies on a limited number of astral MTs engaging cortical Dynein motors. Consequently, decreasing MT numbers generates reduced opposing forces (cortical pushing) that fail to counteract pulling events, leading to a more apical spindle position. Finally, experiments aimed to invert the position (by inhibiting Kinesin-1) of the high- and low-nucleating centrosomes relative to the apical cortex, where the pulling machinery resides, led to two interesting observations. First, NSCs harboring the inverted centrosome configuration, with high nucleating centrosome close to the basal cortex, displayed their spindle positioned toward the apical cortex. Second, this behavior is not due to the loss of Kinesin-1 activity, as the majority of *Khc* mutant NSCs retaining the daughter (high nucleating) centrosome at the apical cortex displayed a basal spindle position similar to control cells.

Our data reveal that final NSC spindle position is dominantly governed by the daughter centrosome that harbors the highest MT nucleation potential.

### Microtubules pushing against the cortex play a dominant role for spindle positioning

The spindle is always positioned away from the daughter centrosome that harbors the highest MT nucleation potential. Additionally, the observation that MT plus ends polymerize against the cell cortex prompted us to propose that asymmetric MT pushing forces could mediate the basal spindle positioning in NSCs. Such MT-based pushing mechanisms against organelles or the cell cortex have been described using *in vitro* reconstituted systems (Dogterom and Yurke, 1997; Holy et al., 1997; Hotani and Miyamoto, 1990) but also *in vivo* where they are sufficient to center the nuclei and spindles in yeast cells and in moving the nucleus toward the anterior cortex in *Drosophila* oocytes (Daga et al., 2006; Tissot et al., 2017; Tolić-Nørrelykke et al., 2004; Tran et al., 2001; Zhao et al., 2012). We experimentally revealed these pushing forces following laser ablation experiments. Ablation of the daughter/apical centrosome triggered displacement of the spindle toward the apical cortex. Conversely, ablation of the mother/basal centrosome triggered displacement of the spindle toward the basal cortex. Altogether, these indicate that pushing MTs are present at both poles in NSCs. Mathematical modeling or *in vivo* biophysical investigations have proposed that these forces may be essential players in spindle positioning in several cell types (Bouvrais et al., 2021; Pécréaux et al., 2016; Yeltokov et al., 2026). We further challenged the importance of cortical pushing forces using 3D agent-centered simulations, namely Cytosim. In these simulations, we modelled apical and basal centrosomes as MT asters with high and low MT counts growing toward the cell periphery, respectively. Astral MTs, upon reaching the cell cortex, generate pushing forces, while the apical pulling force was set as constant on the apical aster. In the presence of low apical pulling force, the asymmetry in MT pushing between the apical and basal cortex was sufficient to recapitulate the basal spindle positioning in wild-type NSCs. Moreover, these simulations, when compared to the corresponding experimental conditions, produced remarkably similar final spindle positioning, when spindles were designed to harbor inverted centrosome configurations or symmetric centrosomes (with low or high centrosome activity).

Our data also shed light on the strength of the apical pulling forces that position the spindle along the apico-basal axis in NSCs. Mud and Pins anchor cortical Dynein at the apical cortex, and *mud* and *pins* mutant NSCs accordingly exhibit spindle orientation defects along this axis (Bowman et al., 2006; Cabernard and Doe, 2009; Gonzalez, 2013; Izumi et al., 2006). Apical Dynein adaptors are thus essential for spindle orientation. Yet they do not alone dictate spindle position in wild-type NSCs: the spindle is invariably displaced basally despite the presence of apical cortical Dynein, and this basal bias is further enhanced when apical pulling is reduced (*Pins* RNAi). Therefore, while the abundant literature has shown that apical pulling is necessary for orientation, our work demonstrates that these forces act in coordination with the apical pushing force exerted by the high nucleating daughter centrosome during spindle positioning. Our 3D simulations support this conclusion: below a certain force threshold, apical pulling cannot draw the spindle apically. To fully understand how cortical pulling and pushing jointly orient the spindle, a more detailed model will be required, one describing pulling events at the scale of individual MTs within the apical cortical crescent.

### A function for centrosome identity in the adaptation of pulling and pushing MT-based mechanisms to cell division type

Our findings uncover a functional role for the pronounced centrosome asymmetry of NSCs. We show that centrosomes retain distinct MT nucleation capacities, inherited from their mother–daughter identity, which generate asymmetric pushing forces that position the mitotic spindle basally during metaphase. This positioning is critical, as it determines the location of the spindle midzone during anaphase and contributes to cleavage site positioning. Consistently, NSCs with abnormally apical spindles give rise to smaller NSCs and larger GMCs. Thus, centrosome-dependent asymmetric pushing forces provide a mechanism linking centrosome identity to spindle positioning and ultimately to the preservation of cell size asymmetry during division.

Together, our findings support a model in which spindle positioning emerges from the balance between opposing MT-dependent forces. While apical Dynein-dependent pulling is necessary for spindle orientation, asymmetric pushing forces generated by astral MTs are essential to maintain the spindle in a basal position. Importantly, our work identifies centrosome-intrinsic differences in MT nucleation as a mechanism that establishes the pushing force asymmetry. These findings complement and extend the recent work of (Yeltokov et al., 2026), highlighting the importance of the interplay between pushing and pulling forces in spindle positioning. By linking this mechanical balance to cleavage site positioning and daughter cell size asymmetry, our study provides a mechanistic framework for understanding how centrosome identity contributes to robust asymmetric cell division, a process whose accurate regulation is essential for tissue homeostasis in the Drosophila central nervous system (Boyer et al., 2026; Delgado et al., 2025).

MT-based pushing forces have long been considered to play a limited role in spindle positioning in most animal cells. Yet, NSCs are relatively small cells (∼12 µm in diameter), with centrosomes positioned close to the cortex, a geometry particularly favorable for MT-dependent pushing. Given the prevalence of small cells in developing tissues, our findings raise the intriguing possibility that MT-based pushing forces may contribute to spindle positioning far more broadly than currently appreciated. More generally, they suggest that intrinsic differences in centrosomal MT nucleation capacity may represent a mechanism by which cells translate centrosome asymmetry into spatial control of cell division.

## Acknowledgments

We thank Timothy Megraw, Cayetano Gonzalez, Paul Conduit, Bill Chia, Anne Ephrussi, Antoine Guichet, and Xin Liang for their kind gift of reagents and Romain Gibeaux for the critical reading of the manuscript. We thank the Bloomington and the Vienna Drosophila RNAi Center for *Drosophila* strains. We acknowledge Stéphanie Dutertre, and Xavier Pinson of the Microscopy Rennes Imaging Center (MRIC, BIOSIT, Biogenouest) for assistance in microscopy. MRIC is member of the national infrastructure France-BioImaging supported by Agence Nationale de la Recherche (ANR-24-INBS-0005 FBI BIOGEN). This project was provided with computing HPC and storage resources by GENCI at TGCC thanks to the grant 2026-<u>A0200315791</u> on the supercomputer Joliot Curie’s ROME partition. IN is a fellow from the Ligue Nationale de la Recherche Contre le Cancer. RG and JP are supported by the Agence Nationale de la Recherche (ANR-24-CE13-2394).

## Methods

### Fly stocks and genetics Fly strains

UAS-Klp10A flies were supplied by C. Dahmann (Max Planck Institute, Germany) (Widmann and Dahmann, 2009). 69B-Gal4 (BDSC 1774), Insc-Gal4 (BDSC 8751), UAS-mCherry-a-tubulin (BDSC 25774 and BDSC 25773) were from the Bloomington Stock Center. The 69B-Gal4 fly stock was used to drive transgene expression in the fly NSCs.

The Insc-Gal4 strain was used to drive over-expression of Klp10A, and mCherry-α-tubulin transgenes in the central brain at 25°C. UAS-Cnb-RNAi (GD28651), UAS-Khc-RNAi (GD12278), UAS-Pins-RNAi (GD11737) lines were obtained from the VDRC.

Eb1-GFP flies (Sun et al., 2021) were kindly given to us by X. Liang (Tsinghua-Peking Joint Center for Life Science, School of Life Sciences, Tsinghua University, Beijing, China). UAS-Cnb-PACT-YFP flies (Januschke et al., 2013) were kindly sent by C. Gonzalez (Institute for Research in Biomedicine Barcelona, Spain), Cnn-sfGFP flies (Mukherjee et al., 2020) were kindly given to us by P. Conduit (Institut Jacques Monod, Paris, France). UAS-GFP-Pon-LD flies (Wang et al., 2007) were kindly given to us by W. Chia (Department of Biological Sciences, National University of Singapore, Singapore). *Khc^63^* (Djagaeva et al., 2012) and *Khc^27^* flies were kindly given to us by A. Ephrussi (European Molecular Biology Laboratory, Heidelberg, Germany,) and A. Guichet (Institut Jacques Monod, Paris France) respectively.

### Antibodies

The following antibodies and concentrations were used in this study: Rabbit anti-PKCξ (C-20, 1:200) was obtained from Santa Cruz Technology. Monoclonal mouse anti-alpha Tubulin (clone DM1A, T2199; 1:200) was obtained from Millipore. Rabbit polyclonal anti-Cnn (Megraw et al., 1999) antibody (1/1000) was kindly given to us by Tim Megraw (Florida State University, USA, Tallahassee). Rat monoclonal anti-Miranda antibody (ab197788, 1:100) was obtained from Abcam. Secondary antibodies were labeled with Alexa Fluor-conjugated (1:200) each obtained from Life Technologies.

### Cell imaging

Third-instar larval brains were dissected in Schneider’s Drosophila medium supplemented with 10% FCS. Isolated brains were loaded and mounted on stainless steel slides, and the preparations were sealed with mineral oil (Sigma-Aldrich) as previously described (Gallaud et al., 2014). Images were acquired with a spinning disk system consisting of a DMi8 microscope (Leica) equipped with a 63X (1.4 N.A.) oil objective, a CSU-X1 spinning disk unit (Yokogawa) and an Evolve EMCCD camera (Photometrics). The microscope was controlled by the Inscoper Imaging Suite and the dedicated software (Inscoper).

For immunofluorescence quantitative analyses, brains were dissected, fixed and incubated with primary and secondary antibodies as described before (Berisha et al., 2024). Fixed immunostained mitotic NSCs were then imaged using a Leica SP8 confocal microscope. Data were processed using ImageJ (Rueden et al., 2017).

### Laser ablation

Laser ablations were performed using a Mai Tai two-photon laser system (Spectra Physics) operating at a wavelength of 800 nm. The laser was coupled to a Zeiss LSM 880 confocal microscope equipped with a Plan-Apochromat 63×/1.4 NA Oil DIC M27 objective and controlled using ZEN Black software (Zeiss). Imaging was performed with the microscope stage maintained at 25°C.Time-lapse images were acquired as z-stacks consisting of seven optical sections spaced 1.5 µm apart and collected every 4 s for 4–5 min. Imaging was performed using 561 nm (1.2% laser power) and 488 nm (0.8% laser power) excitation lasers, with a detector gain of 850, a frame time of 157.29 ms, an image resolution of 256 × 256 pixels, and a pixel size of 87.8 nm.

Laser ablation was performed within a circular region of interest (50-pixel diameter) centered on the centrosome to be eliminated. Ablation was induced between the acquisition of the second and third time points using the Mai Tai laser at 60% power with three iterations. For these experiments, flies expressed mCherry-Tubulin as a mitotic marker, GFP-Pon-LD as a polarity marker, and endogenously tagged Cnn-sfGFP as a centrosome marker. To ensure that centrosome ablation was performed at comparable mitotic stages, only neural stem cells that had undergone nuclear envelope breakdown (NEBD) and entered anaphase at least 1 min 30 s after laser ablation were included in the analysis.

### Analysis of centrosome displacement after opposite centrosome ablation

To analyze centrosome dynamics, a cell-specific coordinate system was defined for each series using the positions of the apical and basal centrosomes at Frame 1. The inter-centrosomal axis was established as the primary reference direction. The unit vector **û** along this axis was calculated as:

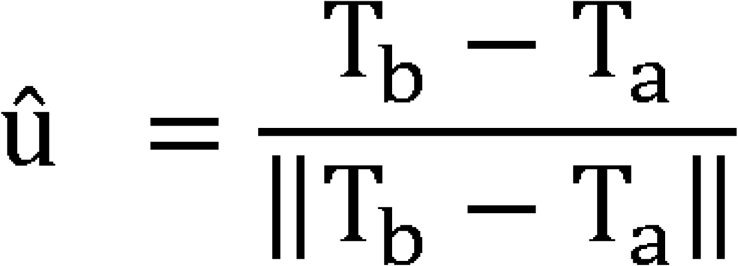

where **T_a_** and **T_b_** are the positions of the apical and basal centrosomes, respectively, and ||·|| denotes the Euclidean norm. For each centrosome, the displacement vector ***d⃗*** between Frames 1 and 2 was decomposed into parallel relative to **û.** The parallel component, **d_par**, was computed as the scalar projection of ***d⃗*** onto **û**:

**d_par** = ***d*** · **û**

This framework enabled the quantification of centrosome movements along the apico-basal axis, facilitating comparisons across cells

### Live and fixed-cell imaging analysis

Image analysis was performed using Fiji/ImageJ software (Rueden et al., 2017) to quantify fluorescence intensities, measure cortex-to-centrosome distances and mitotic spindle lengths, and determine the diameters of NBs and GMCs following cell division.

### Spindle positioning index

Spindle position was quantified as the ratio of the distance between the basal cortex and the basal centrosome (B) to the distance between the apical cortex and the apical centrosome (A). The B/A ratio was log₂-transformed to facilitate visualization. A positive log₂(B/A) values indicates an apical spindle positioning, whereas a negative value indicates a basal spindle positioning. A log₂(B/A) value of 0 corresponds to a spindle positioned at the cell equator (Figure S1).

### Quantification of Cnn fluorescence intensity in fixed and live cells

Cnn fluorescence was quantified using Cnn-sfGFP in live cells and anti-Cnn immunostaining in fixed cells. For both conditions, sum-intensity projections (“Sum Slices”) were generated from Z-stacks. Total fluorescence intensity at each spindle pole was measured using identically sized regions of interest (ROIs) for both centrosomes within the same cell. Background fluorescence was measured in the cytoplasm using a ROI of the same size on the corresponding projection and subtracted from each centrosomal measurement, as previously described (Gallaud et al., 2022).

### Eb1 comets quantification

Z-series were acquired at 1 μm intervals using an LSM 880 confocal microscope equipped with Airyscan (Zeiss) during mitosis. Images were processed using ZEN software. The mean number of EB1 comets nucleated by the basal and apical centrosomes during prometaphase and oriented toward the cell cortex was manually quantified in a single optical section for *n* >10 cells. The number of comets was then extrapolated to the entire 3D volume.

### Simulation

We conducted 3D simulations of the *Drosophila* neuroblast mitotic division using a modified version of Cytosim (Nedelec and Foethke, 2007), which allows multiple fiber types to emanate from a given aster and to apply external forces on solids, fibers or spheres (Pecreaux, 2024). The simulations focused on spindle positioning and the astral microtubule network. Therefore, the spindle was modelled as static: it comprises two disks solidly coupled, mimicking metaphasic sister chromatids, and non-dynamical MTs; spindle length was set to the average *in vivo* values (Figure 1C). The parameters used in the simulations (Table S1) were mostly obtained from literature and are consistent with (Yeltokov et al., 2026). The simulation files are provided as supplemental files S1 to S5 with an exemplar video for each (Videos S3-S7). In further detail, centrosomes were simulated as asters (apical at the top, basal at the bottom) and two types of fibers emanated from each centrosome, spindle- and astral-MTs. A constant external force pointing toward the apical side was applied at the apical centrosome in all simulations to mimic apical pulling. Force magnitude was 5 pN in simulations altering the centrosomal nucleation capacity; and ranged from 0 and 50 pN, when testing the role of pulling force. Astral MTs were defined as fibers with a flexural rigidity of 15 pN.µm^2^, consistent with *in vivo* measurements (Hawkins et al., 2010), with no dynamics at their minus ends. Their plus ends displayed classic dynamic instability. In the wild type simulation, the numbers of MTs nucleated by the apical and basal centrosomes were 123 and 83, respectively. These numbers were extrapolated from the 2D measurements (Figure S3B), considering a focal plane width of 430 nm (Airyscan microscope) and the ability to detect MTs nucleated at the centrosome when longer than 0.75 µm. In the inverted-nucleation simulation, values were swapped. For simulations with symmetric and low (or high) nucleation, MTs numbers were estimated assuming that the number of nucleated microtubules scales linearly with centrosomal Cnn intensity (Figure 2C): 33 for low nucleation and 103 for high nucleation. Astral microtubule dynamics were set according to the literature (Dogterom and Yurke, 1997; Métivier et al., 2020; Rogers et al., 2002).

## Statistical analysis

Statistical analyses were performed using GraphPad Prism 10 (GraphPad Software). Differences between groups were assessed using either the non-parametric Mann–Whitney U test, Wilcoxon test, or one-way analysis of variance (ANOVA) followed by Tukey’s or Dunnett’s multiple comparisons test, as indicated in figure legends. Statistical significance was defined as follows: ns, *P* > 0.05; *, *P* ≤ 0.05; **, *P* ≤ 0.01; ***, *P* ≤ 0.001; and ****, *P* ≤ 0.0001. The statistical tests used are specified in the corresponding figure legends. Each *n* represents an individual neuroblast.

---

**Figure S1.** Scheme for quantification of spindle positioning. **A)** Axial spindle positioning measurements. The apical distance (A) and the basal distance (B) were defined as the distance measured between the daughter/apical centrosome and the apical cortex, and between the mother/basal centrosome and the basal cortex, respectively. B) Schematic showing the expected spindle positioning index (indicated log_2_(B/A) values) of NSCs populations with basal (blue) or apical spindle position. The log_2_ ratio of B/A, hereafter referred as the spindle positioning index was used as a readout of spindle positioning along the polarity axis, with negative values indicating a basal position and positive values indicating an apical position.

**Figure S2.** Analysis of PCM levels in live and fixed cells in *Cnb* RNAi, Cnb-PACT and Klp10A-OE cells. **A)** Selected frames of a NSC from interphase to telophase expressing Cnn-sfGFP, Pon-GFP (green) and mCherry-α-Tubulin (purple). The daughter centrosome (D, green arrow), associated with the apical cortex, retains Cnn-sfGFP and MT nucleation during interphase, while the mother centrosome (M, blue arrow) starts to recruit Cnn-sfGFP and nucleate MTs approximately 6 minutes before NEBD. Scale bar is 10 μm. **B)** Paired quantification (± sd) of Cnn-sfGFP levels between the daughter/apical (green) and mother basal (blue) centrosomes during NEBD (apical: 27100 ± 9697, *n* = 14; basal: 12269 ± 3068, *n* = 14), prometaphase (apical: 28607 ± 18785, *n* = 14; basal: 11686 ± 10006, *n* = 14), metaphase (apical: 23070 ± 8675, *n* = 14; basal: 9275± 3636, *n* = 14), anaphase (apical: 22714 ± 7572, *n* = 14; basal: 10792 ± 2923, *n* = 14), and telophase (apical: 24477± 13589, *n* = 14; basal: 10759 ± 3930, *n* = 14). ***, *P* ≤ 0.001; ****, *P* ≤ 0.0001 (Mann-Whitney test). The daughter and the corresponding mother centrosome are connected by straight lines. **C)** Quantification of centrosomal Cnn-SfGFP asymmetry throughout mitosis, expressed as the ratio of Cnn-sfGFP fluorescence intensity at the daughter (apical) centrosome to that at the mother (basal) centrosome. Each dot represents the ratio calculated for an individual NSC at the indicated mitotic stage. Ratios greater than 1 indicate apical Cnn enrichment, whereas a ratio of 1 indicates equal Cnn-sfGFP intensity at both centrosomes. NEBD (2.24 ± 0.65, *n =* 14), prometaphase (2.70 ± 0.64, *n* = 14), metaphase (2.75 ± 0.70, *n* = 14), anaphase (2.13± 0.45, *n* = 14), and telophase (2.25± 0.55, *n* = 14). **D)** Dot plot showing the mean (± sd) of the Cnn centrosomal protein ratio between spindle poles in control, Cnb-PACT, *Cnb* RNAi, and Klp10A-OE metaphase-fixed NSCs. Control: 2.65 ± 0.77, *n* = 16; Cnb-PACT: 1.05 ± 0.09, *n* = 14; *Cnb* RNAi: 1.02 ± 0.20, *n* = 16; Klp10A-OE: 2.10 ± 0.65, *n* = 15. Ratios > 1 indicate apical Cnn enrichment, whereas a ratio of 1 indicates symmetric Cnn distribution between spindle poles. ****, *P* ≤ 0.0001 (Mann-Whitney test).

**Figure S3.** Analysis of MT nucleation by the mother and daughter centrosomes during mitosis. **A)** Selected frames of a prometaphase wild-type cell expressing Eb1-GFP. The left panel shows a 5 μm projection (5 planes), the middle and right panels show confocal sections of the daughter/apical centrosome (indicated by green triangles) and the mother/basal centrosome (indicated by tblue triangles) respectively. Scale bar is 2 μm. See also Video S2. **B)** Quantification of the mean number of Eb1-GFP comets number (± sd) nucleated by each centrosome and pointing toward the cell cortex in a single confocal section. Daughter centrosome: 33.9 ± 10.8, *n =* 44, mother centrosome 22.5 ± 7.5, *n =* 44. ****, *P* ≤ 0.0001., Wilcoxon test.

**Figure S4.** Simulated NSC spindle positioning with increasing apical pulling. A-A’’’) Representative images of the final simulated spindle position in mitotic NSCs under four experimental conditions, in which 0, 15, 30 and 50 pN as pulling forces were applied. Upper panels and lower panels show simulated 3D projections and single optical sections, respectively. **B**) Violin plot showing the mean log_2_(B/A) measurements (spindle positioning index) of the final NSC spindle positions (± sd) subjected to increasing apical pulling. 0 pN: -0.47 ± 0.13, 5 pN: -0.24 ± 0.14, 10 pN: 0.07 ± 0.14, 15 pN: 0.10 ± 0.16, 20 pN: 0.28 ± 0.16, 25 pN: 0.46 ± 0.16, 30 pN: 0.60 ± 0.15, 35 pN: 0.78 ± 0.18, 40 pN: 0.91 ± 0.15, 45 pN: 1.05 ± 0.15, 50 pN: 1.17 ± 0.13. For each simulated condition, *n* = 100

**Video S1. NSC expressing Cnn-SfGFP, GFP-Pon and Ch-tubulin during cell division.**The GFP-tagged proteins are shown in green (left) and monochrome (right). Scale bar is 10 μm. Time is min:sec. Corresponding to Figure S2.

**Video S2. NSC expressing Eb1-tdGFP during metaphase.** Scale bar is 2 μm. Time is sec. From left to right: 5 μm projection (5 planes), apical/daughter centrosome focal plane, basal/mother centrosome focal plane. Corresponding to Figure S3.

**Video S3. Cytosim simulation of spindle positioning (3D view) corresponding to a wild type NSC with asymmetric pushing forces mediated by astral MTs and subjected to 5 pN of apical pulling.** Corresponding to Figure 5A.

**Video S4. Cytosim simulation of spindle positioning (3D view) corresponding to a NSC with symmetric high number of MTs emanating from the centrosomes, causing strong pushing forces mediated by astral MTs and subjected to 5 pN of apical pulling, similar to Cnb-PACT expression.** Corresponding to Figure 5A’.

**Video S5. Cytosim simulation of spindle positioning (3D view) corresponding to a NSC with symmetric low number of MTs emanating from the centrosomes, causing weak MT-based pushing forces mediated by astral MTs and subjected to 5 pN of apical pulling, similar to *Cnb* RNAi.** Corresponding to Figure 5A’’.

**Video S6. Cytosim simulation of spindle positioning (3D view) corresponding to a NSC with asymmetric but inverted number of MTs emanating from the centrosomes, causing reversed pushing forces mediated by astral MTs, similar to *Khc^63^/Khc^27^* mutants with inverted centrosome configuration.** Corresponding to Figure 5A’’’.

**Video S7. Cytosim simulations of spindle positioning (3D view) corresponding to a wild type NSC with asymmetric number of MTs emanating from the centrosomes and thus pushing forces mediated by astral MTs and subjected to increasing apical pulling forces.** The apical pulling forces (0, 15, 30, 50 pN) are indicated at the top left corner. Corresponding to Figure S4.

## Notes

### Competing Interest Statement

The authors have declared no competing interest.

