## Supplementary figures and images for "Centrosome identity drives asymmetric microtubule-dependent pushing forces, spindle positioning and daughter cell size"

### Figure S1

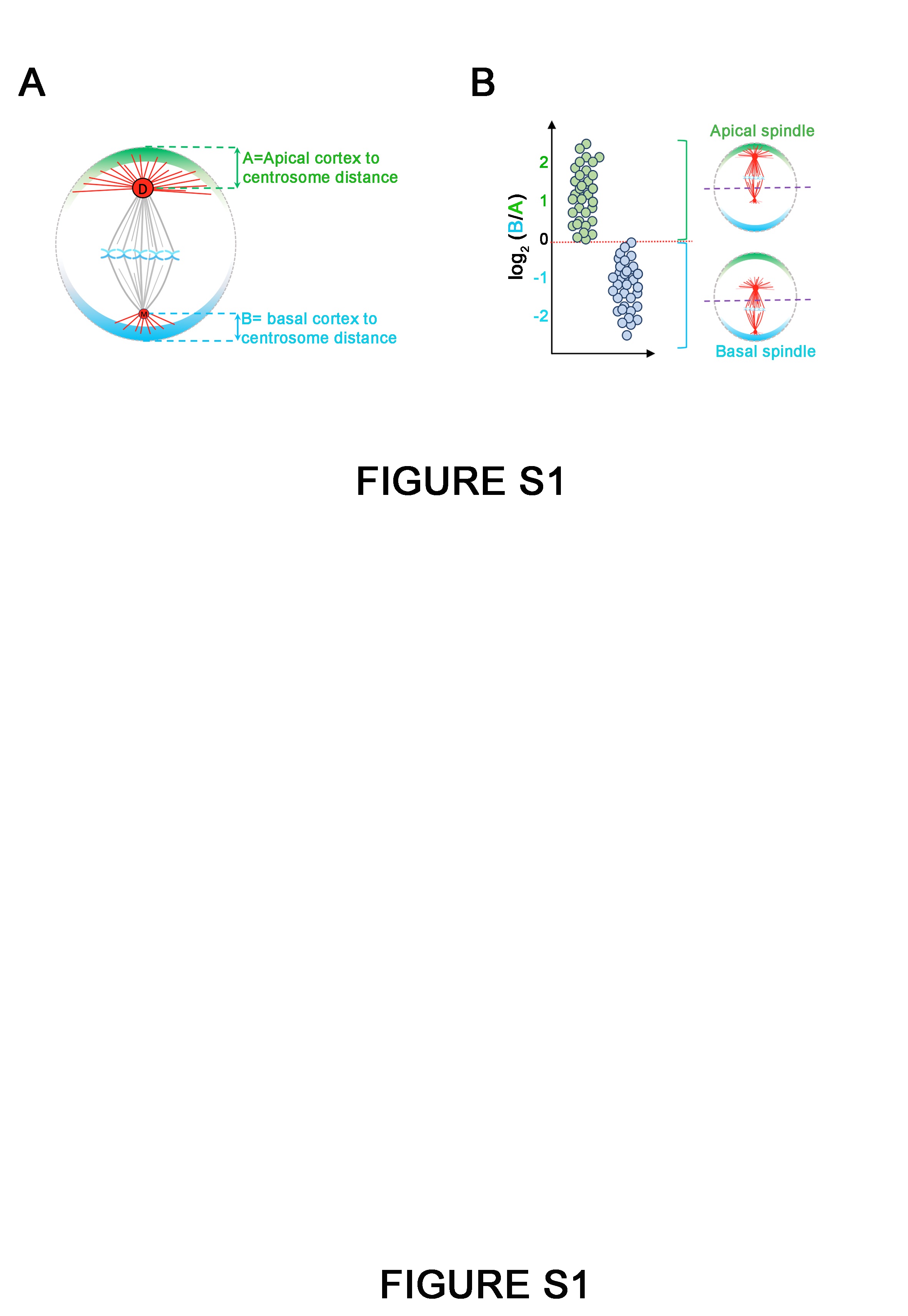

### Figure S2

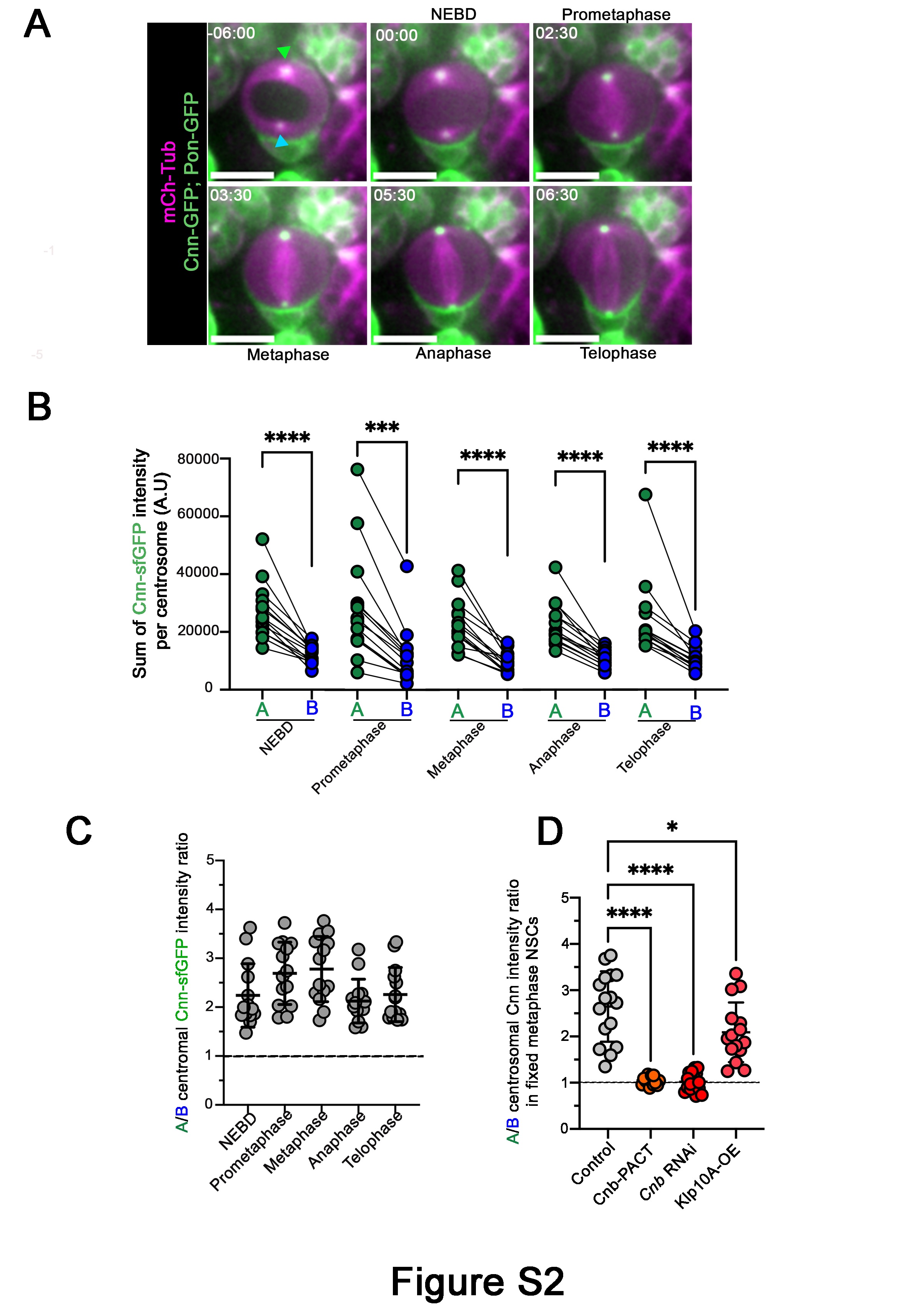

### Figure S3

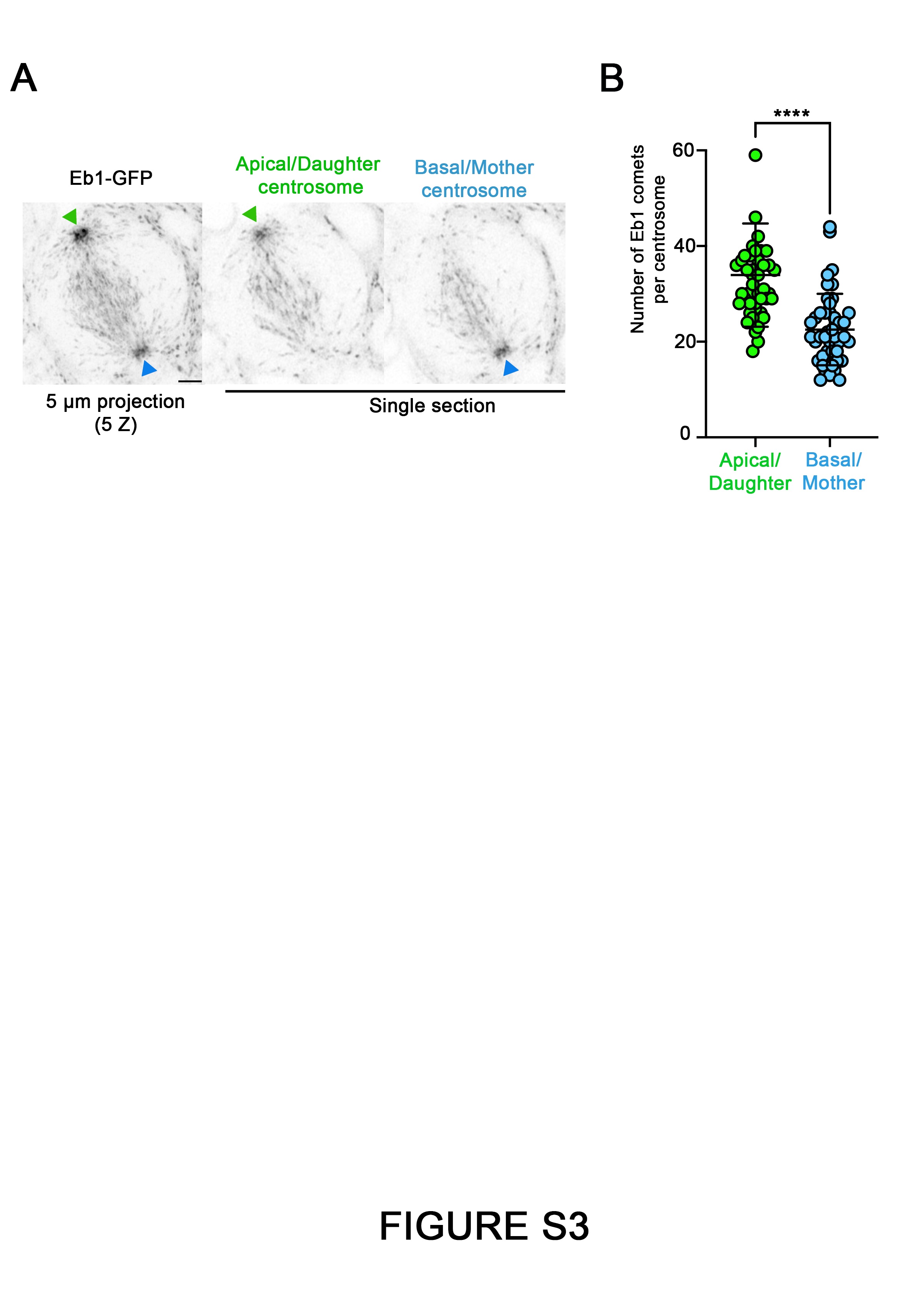

### Figure S4

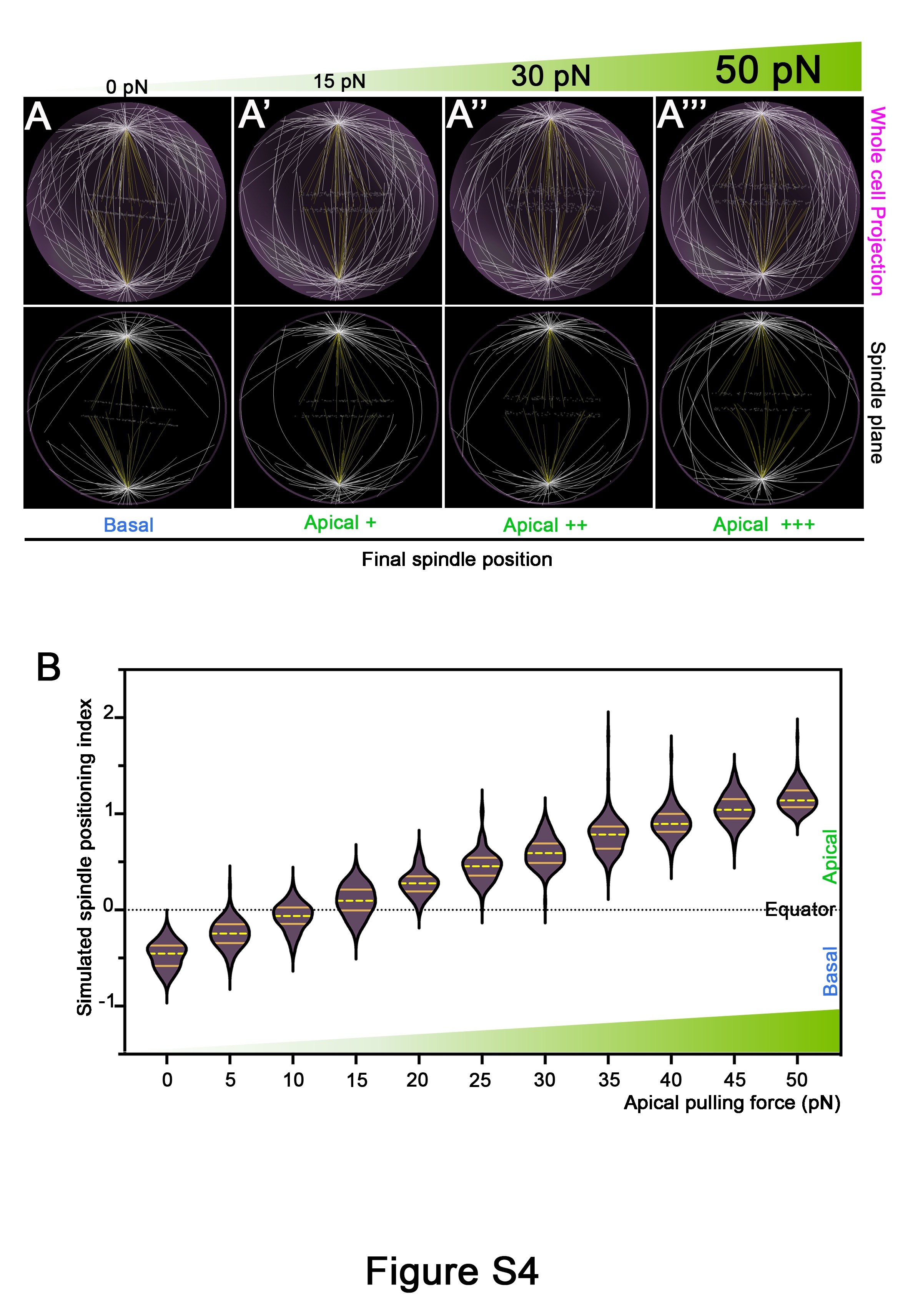
